# Genetical control of 2D pattern and depth of the primordial furrow that codes 3D shape of the rhinoceros beetle horn

**DOI:** 10.1101/2020.05.26.116327

**Authors:** Haruhiko Adachi, Keisuke Matsuda, Teruyuki Niimi, Shigeru Kondo, Hiroki Gotoh

**Author notes:** Corresponding author and address. Ecological Genetics Laboratory, Department of Genomics and Evolutionary Biology, National Institute of Genetics, Mishima, Shizuoka, 411-8540, Japan.

## Abstract

The head horn of the Asian rhinoceros beetle develops as extensively folded primordia before unfurling into its final 3D shape at the pupal molt. The information of the final 3D structure of the beetle horn is encoded in the folding pattern of the developing primordia. However, the developmental mechanism underlying epithelial folding of the primordia is unknown. In this study, we addressed this gap in our understanding of the developmental patterning of the 3D horn shape of beetles by focusing on the formation of surficial furrows that become the bifurcated 3D shape of the horn. By gene knockdown screening via RNAi, we found that knockdown of the gene *Notch* disturbed overall horn primordia furrow depth without affecting 2D furrow pattern. In contrast, knockdown of *CyclinE* altered 2D horn primordia furrow pattern without affecting furrow depth. From these results, depth and 2D pattern of primordial surficial furrow are likely to be regulated independently during the development and both of change can alter the final 3D shape.

**Author Summary:** In insects, some large structure is made under the old exoskeleton before the molting. Long horn of rhino-beetle is one of extreme cases. The beetle horn is compactly packed as furrowed primordia under the larval exoskeleton. At molting, the primordia is extended to form its final 3D horn shape as blowing up furrows like a balloon. This transformation from primordia to final horn does not required any living cell activities. Thus, characteristics of furrows of primordia actually determine the final 3D shape. However, molecular mechanisms and genetic basis of furrow formation is not well understood not only in beetle horn but also in any other insects. In this study, by using beetle horn as a model, we addressed what kind of genetic factors are contributed to primordial furrow formation. By gene knockdown screening, we found that knockdown of the gene *Notch* disturbed primordial furrow depth without affecting 2D furrow pattern. In contrast, knockdown of *CyclinE* altered 2D furrow pattern without affecting furrow depth. In both case, final horn shapes were disturbed. From these results, we concluded that both of the depth and 2D pattern of primordial furrow can contribute final shape, but their development is controlled independently.

## Introduction

Insect body is covered with hard exoskeleton structure, so which size is increased via molting. Prior to molt, they form a new furrowed cell sheet under the old exoskeleton, and the furrow is extended after molting. In many hemimetabolous insects, the shape after molting is similar to before one. However, in some holometabolous insects obtain their large 3D organs through the molting, like the wing, the leg, and the horn. In *Drosophila* wing and legs, after eversion of the primordia, further expansion and elongation process takes over seven hours and involves biological factors (e.g. cell division, cell movement and cell deformation) [1, 2]. On the other hand, Other insect traits, such as the exaggerated horn of the beetles, also develop under the larval cuticle, but the beetle horn appears in a relatively short period of time (within two hours) at pupation^3^. The horn primordia consist of a furrowed epithelial cell sheet with cuticle located under the larval head cuticle. At molting, the primordia is extended to form its final 3D shape as blowing up furrows like a balloon. This transformation from furrowed primordia to final horn does not required any living cell activities [3]. This indicates that the information for the final 3D structure of the beetle horn is patterned within the primordia by this time. The structure of the horn primordia, which is comprised of both the depth and pattern of the surficial furrow, determines the final 3D shape of horn. The primordial structure consists of mushroom-like macro structure of the entire primordia (hereafter “macro structure”) and surface micro furrows (hereafter “micro furrows”) [3, 4] (MovieS1, Fig. S1). Both macro structure and micro furrows contribute to the final 3D shape of the beetle horn [3, 4]. In our previous study, we showed that anisotropy of cell division contributes to macro structure formation, especially in the stalk region of the mushroom-like structure [4]. What remains unknowns is the morphological, molecular and cytological properties of the micro furrows which are not well understood. In this study, we investigated the factors determining the property of micro furrows in the developing beetle head horn primordia, and linked them to the final 3D horn structure. We focused on the cap region of the mushroom-like primordia because this region is crucial for the tetra-furcated horn shape [5]. The cap region in the primordia has four morphological parameters as follows: the (1) size of macro structure, (2) depth, (3) density and (4) the 2D pattern of the micro furrows. These parameters directly determine the 3D shape of the horn. Our objective was to study how these four parameters are regulated. First, we focused on size-dependent differences of the beetle horn to clarify which parameters are dependent on body size and which is not. Second, we screened genes that produced RNAi-induced abnormal head horn phenotypes to find genes which affect the primordial parameter(s) accounting for final horn shape. As results, we found that primordial parameters are controlled independently by different factors and gene function.

## Results and Discussion

### The depth of the furrow and 2D furrow pattern is independent of body size

Pupal beetle horns are known to differ in length depending on body size, mainly due to growth conditions [6-9] (Fig. 1A). The size of the tetra-furcated tip region of horn also had similar body size dependent differences (Fig. S2). We investigated how the macro structure and micro furrows of the primordia contributed to that size difference of the tip region and whether those parameters fluctuate simultaneously or independently. We analyzed the depth and density of furrows and mushroom-like macro structure size in different size primordia because these factors can change the horn size (Fig. S3). We found that the depth of furrows was constant regardless of the body size (Fig. 1C and D).

**Figure 1.**
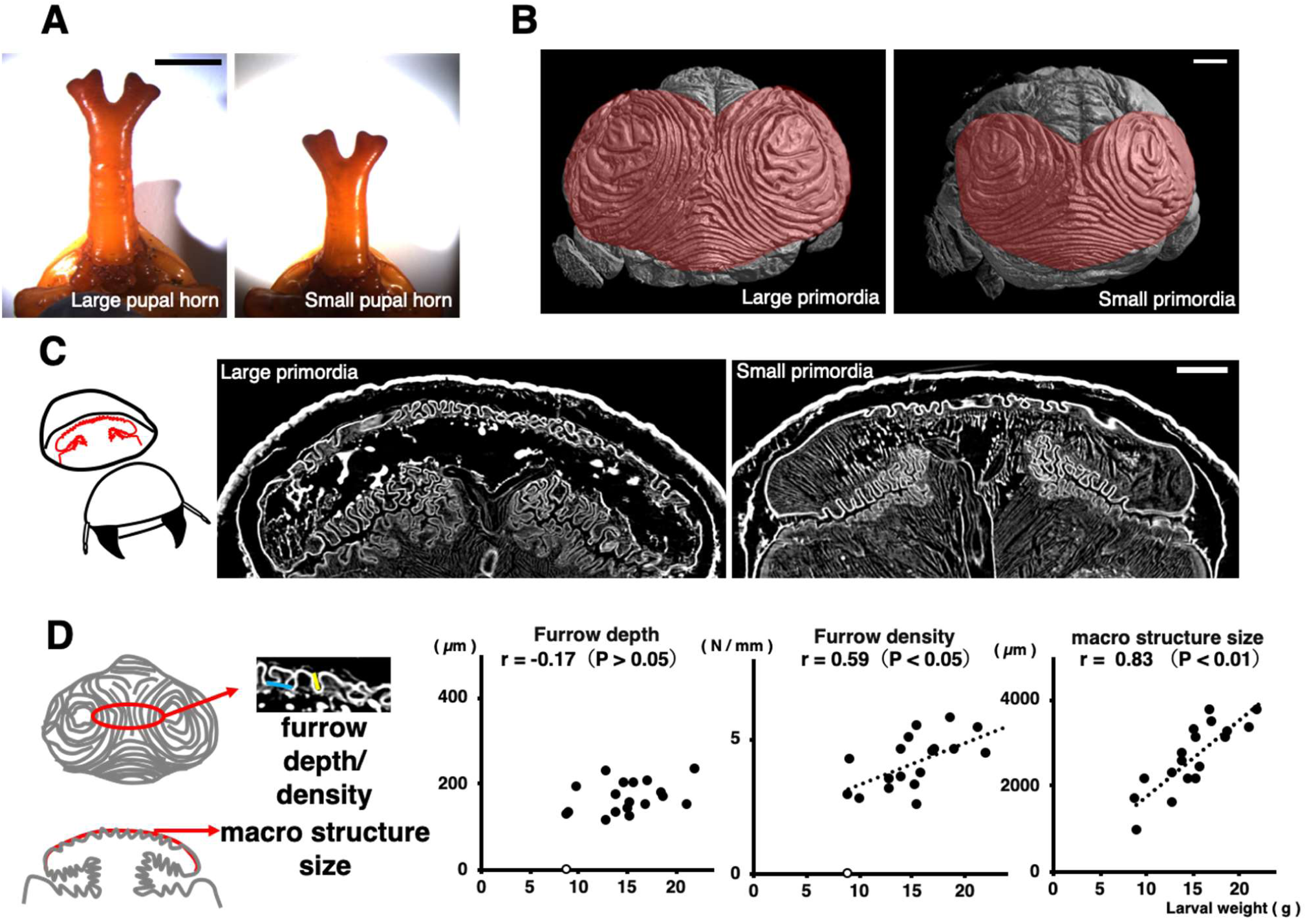
Relationship between body size and the morphology of the primordia. (A) The horn of pupae with different body sizes (left: 18 g, right: 14 g). Larger beetle has longer horn. (B) The primordia of the horn from different body sizes (left: 18 g, right: 14 g). Both of the primordia have similar 2D furrow patterns (concentric furrow patterns and stripe furrows between them), while the overall size of primordia is different. The cap (top) regions of the primordia are indicated in red color. The images were acquired by µCT scanning. (C) Virtual frontal section images of horn primordia via µCT scanning. The mushroom-like macro structure can be recognized regardless of body size. (D) Relationship between body size and the macro structure size, the density of the furrows, and the depth of the furrows. For macro structure size the correlation coefficient was 0.83 (p < 0.01), for furrow density the correlation coefficient was 0.59 (p < 0.05), and for furrow depth, the correlation coefficient was not significant. In the smallest beetle (white plot: 8.95 g), obvious furrows could not be detected so it was excluded from the analysis (n = 19). Scale bar is 10 mm for (A), 1 mm for (B) and (C).

The density of furrows was negatively correlated with body size, but its correlation coefficient was moderate (Fig. 1C and D) or did not correlate in some areas (Fig. S4 and S5). On the other hand, the macro structure size of the primordia was strongly correlated with the body size (Fig. 1C-D, Fig. S4). This tendency (constant furrow depth, weak or no correlation of furrow density and strong positive correlation of macro structure) was also observed other regions of horn primordia (Fig. S4 and S5).

We also observed primordial surface 2D furrow patterns among different body sized animals. The surface furrow pattern (a pair of concentric-like furrows pattern) was similarly observed in both small and large larval primordia (Fig. 1B). These data suggested that size-dependent morphological change in horns is not mainly controlled by changing the depth of the primordia but by changing the macro structure size.

### The Notch and CyclinE genes contribute to the control of furrow depth and 2D furrow pattern, respectively

From these observations, it is assumed that the depth and 2D patterns of the furrows are strictly regulated independently of the body size. Hence, we next searched for the gene(s) involved in controlling furrow depth and 2D pattern. Methodology for use of RNAi is well established for beetles, and the horn shape can be changed by specific gene knockdown [9-12]. We screened six candidate genes (*Notch* <*N*>, *CyclinE* <*CycE*>, *dachsous* <*ds*>, *mushroom body defect*<*mud*>, *Optix* <*Optix*>, *Retinal Homeobox* <*rx*>), those genes are already known to play important role in insect organogenesis including beetle horn [4, 11, 13-19], using RNAi to see how they affected the furrows on the horn primordia.

Although we found several differences in primordia development in RNAi animals, we especially focused *N* RNAi and *CycE* RNAi because these two more drastically changed the depth and 2D pattern of furrows (Fig. 2A) than other candidate genes. *N* RNAi decreased the depth of furrows relative to controls (*egfp* RNAi) in almost all measured regions of primordia (Fig. 2B-C, Fig. S7), except for a pair of specific furrows (Fig. 2B red arrowhead). In addition to decreasing furrow depth, macro structure size was increased and the furrow density was decreased in *N* RNAi beetles (Fig. 2C, Fig. S7). We then manually extended the primordia to investigate the influence of furrow depth change on the final horn shape, because none of *N* RNAi larvae survived until pupation. We found that in these manually extended horns the center groove and side groove were missing and the shape of the horn tips were similar to the centroclinal primordial shape (Fig. 2D). On the other hand, *CycE* RNAi changed the 2D furrow pattern (from concentric-like to zigzag-like) (Fig. 2A, Fig. S8) and the pupal horn shape (the center groove became shallower) (Fig. 2E), while the macro structure size, and the density and depth of the furrows were not affected (Fig. 2B-C, Fig. S7).

**Figure 2.**
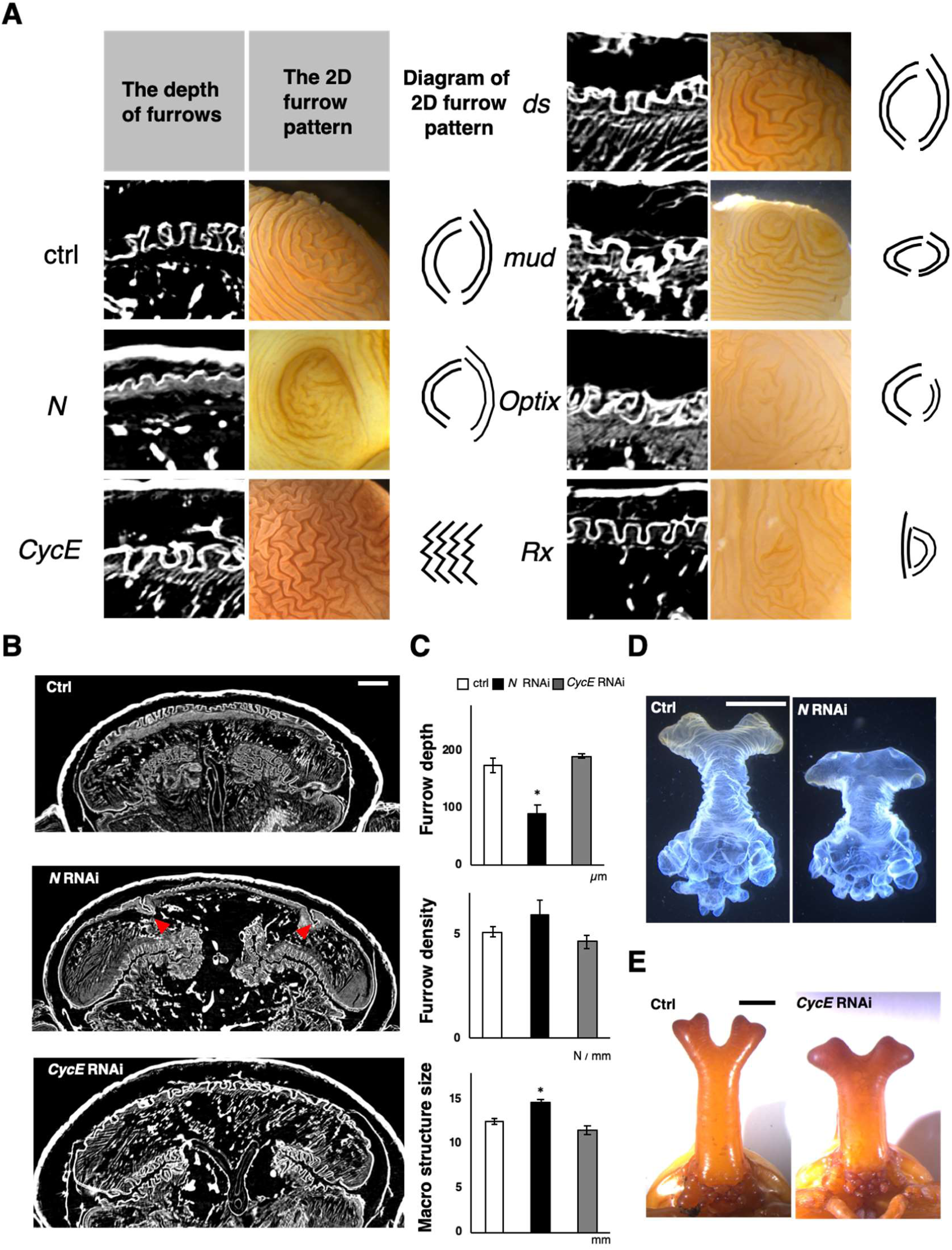
RNAi screening show that the *Notch* and *CyclinE* genes contribute to control of furrow depth and 2D furrow pattern, respectively. (A) Comparison of the depth and surface 2D pattern (concentric-like pattern) of the horn primordia between control and six different RNAi treatments. The 2D furrow patterns inside of the primordial top region were compared. *N* RNAi decreases the depth of the furrow and *CycE* RNAi disturbed the concentric-like 2D furrow pattern. (B) Comparison of the frontal section of the head just before pupation in control and *N* RNAi and *CycE* RNAi. The red arrowheads show the specific deeper furrows detected in *N* RNAi. (C) Quantitative data of the macro structure size and the density and the depth of the furrow (asterisk means P < 0.05) (n = 7, 5, 6 for negative control, *N* RNAi, *CycE* RNAi, respectively; however the furrows were too shallow to measure in one *N* RNAi so that n = 4 for furrow depth and density). (D) Comparison of the final horn shape by extending the primordia artificially, between control and *N* RNAi. (E) Comparison of the pupal horn shape between control and *CycE* RNAi. Scale bar indicates 1 mm for (B), 5 mm for (D) and (E).

Recently, the contribution of Notch signaling to horn formation was also demonstrated in a dung beetle, *Onthophagus taurus* [19]. Our result is consistent with this, and we have found that Notch plays an important role in determining the final horn shape via regulating primordial furrow depth. In addition to the depth of the furrows, the density of the furrows and macro structure size of the primordia were also affected by *N* RNAi. Considering that the macro structure size and the density of the furrow can vary depending on body size, while the furrow depth did not, it is assumed that *N* RNAi has a direct effect on the depth of the furrow (i.e. the furrow density increasing in *N* RNAi likely to be a byproduct of the effect on furrow depth).

As for the relationship between the morphology of the primordia and pupal horn in *N* RNAi, it is suggested that the change of the depth of furrows causes the change of the final horn shape. Also, in *CycE* RNAi, it is presumed that the change of 2D furrow pattern causes the change of the pupal horn shape, because the different 2D furrow pattern can be extended to variable 3D structure [3]. These results strongly indicate that both furrow depth and 2D furrow pattern are important parameters affecting final horn shape, but that they are regulated independently.

### The Notch and CyclinE genes contribute to the control of frequency and localization of mitosis, **<u>respectively</u>**

Next, in order to investigate the relationship between cell division and 2D furrow pattern and furrow depth, we analyzed the orientation, frequency and localization of cell division in the mushroom-shaped cap region (Fig. 3A) among control, *N* RNAi and *CycE* RNAi primordia.

**Figure 3.**
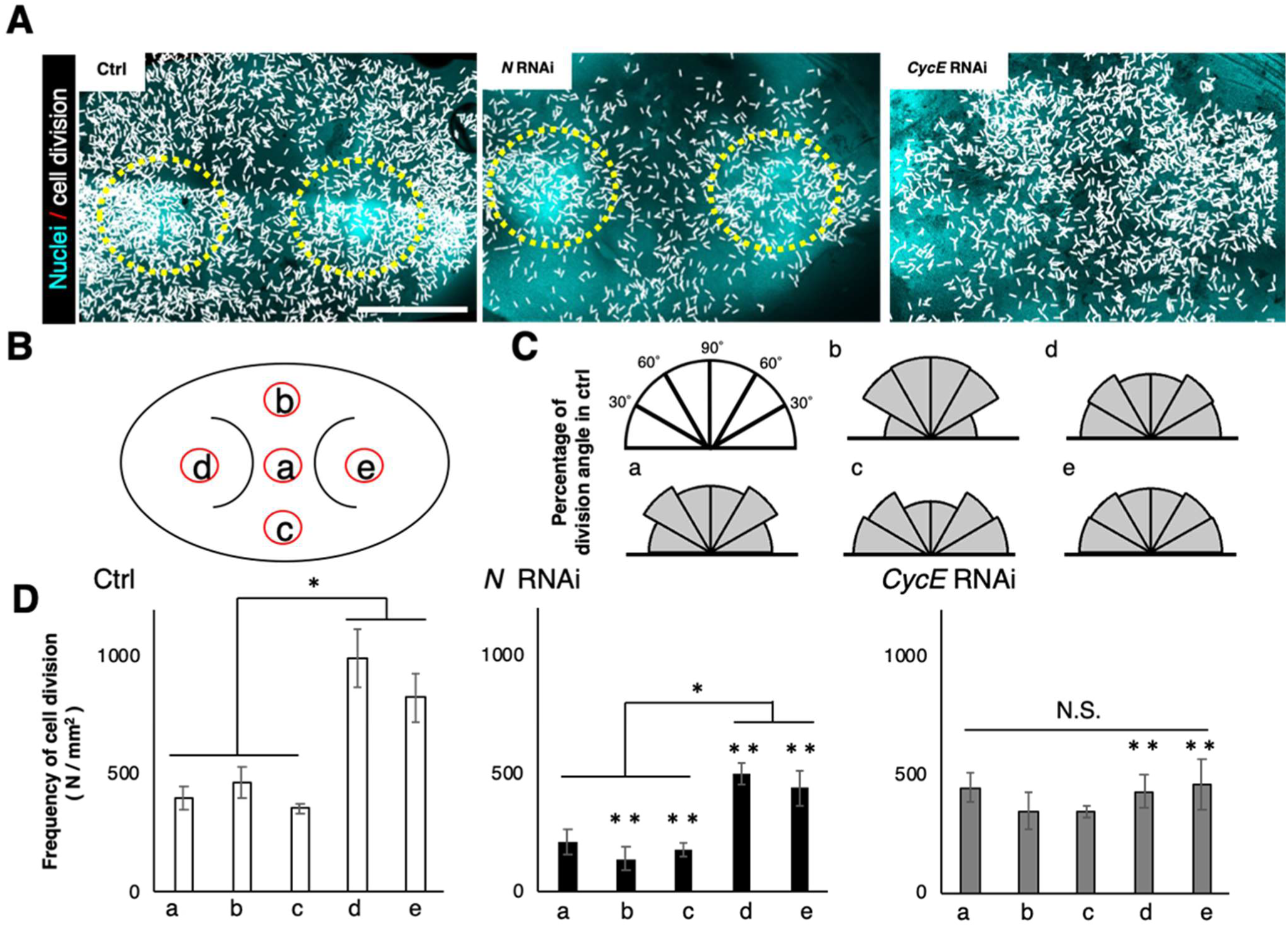
Analysis of cell division of the primordia. (A) Plot of the orientation and the localization of cell division in control and *N* RNAi and *CycE* RNAi. Each white line indicates one dividing cell. Direction of white line shows the orientation of cell division at that point. Yellow circles show the area of high intensity of Hoechst fluorescence. (B) The area of measured cell division. Five regions were determined by using a pair of characteristic crescent shape furrows as landmark. This furrow stably formed as the first furrow in all of the analyzed primordia (Figure 3—figure supplement 1). (C) Plot of the percentage of the orientation of cell division in five areas of the primordia. Significant change of cell division orientation was not detected (n = 5, for negative control). The results for RNAi are shown in Figure 3—figure supplement 1. (D) Quantification of frequency of cell division in five areas of the primordia. In control and *N* RNAi, the frequency of mitosis in the right and left parts were twice as high as in the other parts. *N* RNAi decreased the number of cell divisions in all areas of the primordia. *CycE* RNAi decreased cell division only in both of the side areas of the primordia. Single asterisk (*) indicates significant (P < 0.05) difference between the primordia areas in each RNAi treatment. Double asterisk (**) indicates the statistical significance comparing the same area of primordia between *N* or *CycE* RNAi and negative controls (n = 5, 4, 5 for negative control, *N* RNAi, *CycE* RNAi, respectively). Scale bar indicates 1 mm for (A).

In all of the experimental groups, including the control, there was no clear anisotropy of cell division in any measured region of the primordial cap (Fig. 3B-C, Fig. S9). Thus, region-specific anisotropy of cell division is not likely to be involved in furrow depth control and region-specific 2D furrow patterns. On the other hand, in control primordia, a specific localization pattern of mitosis was observed among the regions (Fig. 3D). The frequency of mitosis in the right and left parts were twice as high as in the other parts (Fig. 3D). In *N* RNAi, total mitosis was decreased in all of the measured regions, while its distribution pattern was not changed (Fig. 3D). In *CycE* RNAi, the specific localization pattern of cell division was disturbed. That is, the frequency of mitosis in the right and left parts of primordia was decreased, which resulted in a uniform distribution of mitosis across the regions (Fig. 3D).

A numbers of studies have reported that Notch contributes to cell proliferation in insect development *(i.e.* labrum of *Tribolium castaneum*, the eye and the wing of *Drosophila melanogaster*) [20-22]. Hence, in beetle horn development, Notch is also assumed to contribute to cell proliferation.

However, the relationship between the frequency of mitosis and the depth of the primordial furrow is still unknown. Notch signaling is also well known to contribute to joint formation in insects[17, 23]. The mechanism of furrow formation may be similar to joint formation because both of them include the folding of epithelial cell sheets [24]. In the future, research about joint formation may provide avenues to further understand furrow formation.

As can be seen from nuclei staining images, the fluorescence intensity of Hoechst was higher in specific areas of control and *N* RNAi horn primordia (Fig. 3A yellow circle), but this fluorescence pattern was not detected in *CycE* RNAi. This means these brighter areas contained more cells in the S/G2 phase of the cell cycle because the fluorescence intensity of Hoechst is dependent on ploidy. The *CycE* gene is involved in the progression of the cell cycle, especially the transition from G1 to S phase [25]. Hence, it assumed that *CycE* RNAi decreased mitosis in a specific area. The region where the frequency of cell division was disturbed in *CycE* RNAi and the region where the 2D furrow pattern (concentric-like pattern) was disturbed in *CycE* RNAi were identical, which suggested that the cell division distribution pattern can affect the 2D furrow pattern. Although the developmental link between the cell division distribution and final 2D furrow pattern is still unknown, one of the possibilities is that specific mechanical stress caused by differences of cell division frequency may determine the direction of furrows.

## Conclusion

In this study, we revealed that morphological parameters (macro structure and micro furrow 2D pattern and depth) in the horn primordia were controlled by different mechanisms. That is, macro structure size is controlled in a body-size dependent manner while micro furrow depth and 2D pattern are not. The depth and 2D pattern of micro furrows are controlled independently via Notch signaling and CyclinE-dependent mechanisms, respectively (Fig. 4). Any one of these morphological parameters can be changed independently, which then alters the final 3D horn shape as well.

**Figure 4.**
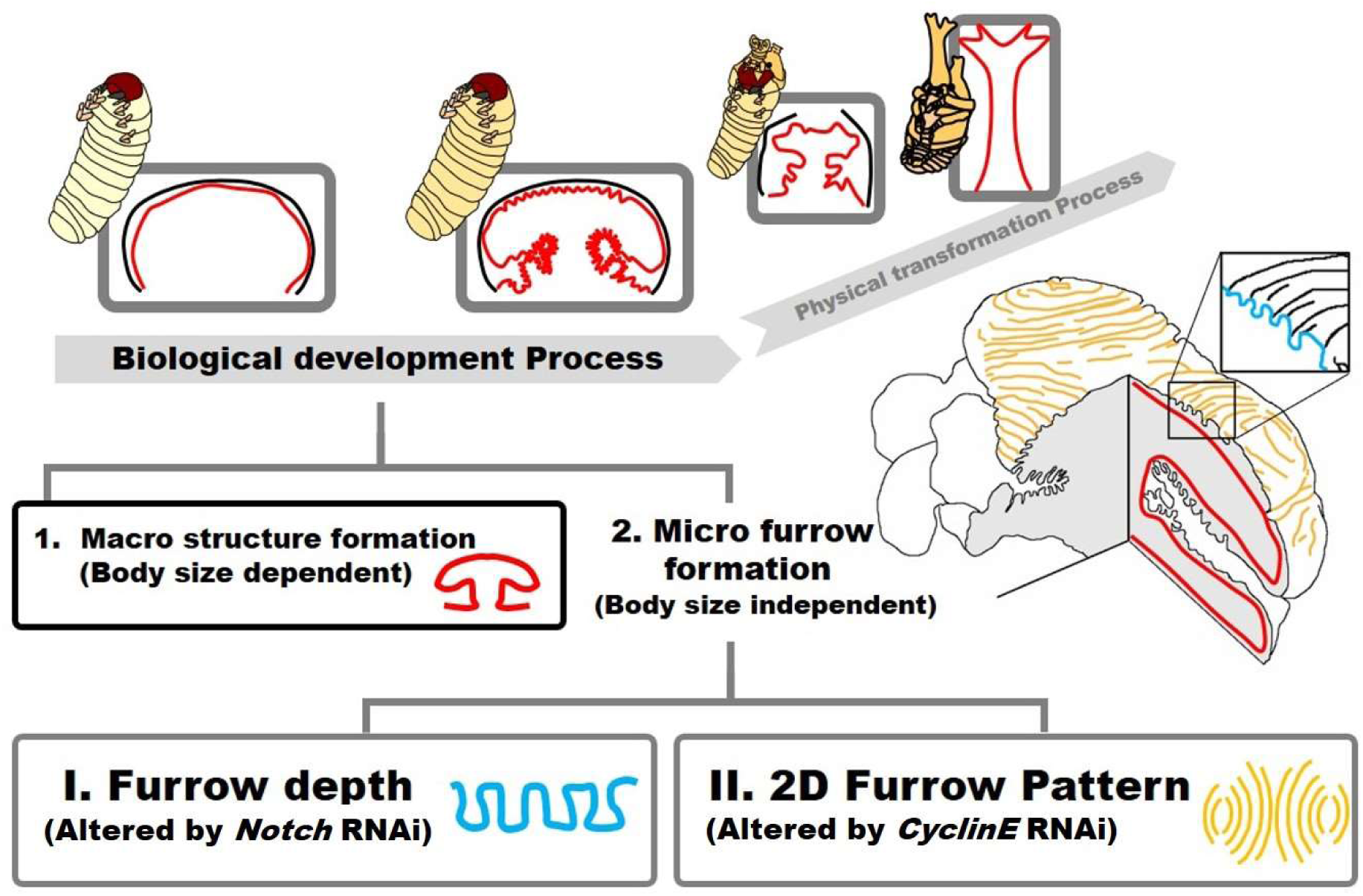
Summary of beetle horn development via “folding and extension”. In the “folding and extension” seen in beetle horn development, the final 3D shape has been already coded when the folding process is done. In this study, we demonstrated that the folding process can be divided into 1) macro structure formation and 2) micro furrow formation. Micro furrow formation can be further divided into depth regulation and 2D pattern formation via Notch and CyclinE control, respectively.

## Material and Methods

### Insects

The beetle larvae were purchased and kept according to our previous studies[3]. Briefly, commercially purchased last instar (third instar) larvae of the Asian rhinoceros beetle *Trypoxylus dichotomus* were kept individually in 1 L or 800 mL plastic bottles filled with rotting wood flakes at 10–15 °C to suspend their development. Larvae were moved to 25 °C to restart their development before the experiments and/or observations. Bottles were checked daily in order to record the initial date of pupal chamber formation. We defined Day 1 as the first day when the pupal cell was clearly recognized. Most male prepupae pupated at Day 8, therefore we used Day 8 horn primordia with brownish color (a sign of sclerotization) as fully developed ones. Pupae and prepupae were weighed before each experiment. We used 7.28 to 18.05 g pupae and 8.95 to 21.30 g prepupae for detecting size-dependent parameters (Fig. 1). For RNAi screening and RNAi-induced horn primordia phenotype analyses, we used a range of pupae from 13.66 to 23.35 g.

### Analysis of pupal horn morphology

Pupal horns were photographed using a digital camera (MZ16FA, Leica, Germany). The inflection point was used as a landmark and the length was measured (Fig. S2) using Fiji (Image J). For the length of the side groove and cap side stalk, both sides (right and left) were measured and averaged.

### Degradation of inner structure of the primordia

The primordia have an inner structure containing muscle, tracheae and body fluid in addition to cuticle. To observe cuticular structure clearly, we degraded the inner tissue by incubating overnight with 10% KOH solution at 60 °C followed by washing with DDW and dehydration with ethanol. The dehydrated sample was soaked with t-butanol and freeze-dried. The sample was used for µCT observation and expansion experiments to observe the final horn shape of notch RNAi.

### µCT scanning

Prepupae were anesthetized on ice and frozen using liquid nitrogen. The frozen samples were truncated by sawtooth. The truncated heads were dried using a freeze-drying system (FZ-2.5, Asahi Life Science, Japan) over 7 hours. Then the dried samples were scanned using a micro-CT scanner (Skyscan1172, Bruker, USA) following the manufacture’s instruction. The X-ray source ranged from 40 kV, and the datasets were acquired at a resolution of 9 μm / pixel. The stacks of transverse sections were reconstructed from primary shadow images using SkyScan software NRecon. From these image stacks, 3D volume-rendered images were constructed using SkyScan software CT Vox.

### Analysis of the horn primordia morphology

The equal-section images were obtained using landmarks such as the points changing furrow direction and inflection points of macro structures (Fig. S6) using SkyScan software DataViewer. Firstly, the length of the cap region of the mushroom-like macro structure of the section images was measured. Then the density of furrows was calculated from the number of furrows per the length of measured region. The average depth of individual furrows was calculated from the depth of all furrows in the measured region. All data were measured using Fiji (Image J). Also, in the bottom region of the mushroom-like macro structure, the same analysis was conducted.

### Nuclei staining

Prepupae were anesthetized on ice before dissection. Amputated prepupal heads were fixed with 4% PFA for 2 days at 4 °C. Dissections were performed under a binocular microscope (SZ61-TRC, Olympus, Tokyo, Japan) Dissected PFA-fixed primordia were further divided into small pieces under a binocular microscope. During this process, the original position within each primordia fragment was recorded. Fragment tissues were washed with PBS three times and incubated at room temperature for 60 minutes with Hoechst 33342 (1:1000, Invitrogen) in blocking solution (1% BSA), then washed with PBS three times, before tissues were mounted on a glass slide and covered with a cover glass. Fluorescent images were observed and recorded with a confocal laser scanning microscope (LSM-780; Carl Zeiss, Jena, Germany).

### Analysis of cell division

Dissected cap area tissue from developing horn primordia at prepupal Day 4 was stained by Hoechst because the timing is just before formation of many micro furrows (Fig. S10) and cell division occurs frequently during this time. All of the dividing cells and the angle of each division of the primordia were measured using Fiji (Image J) [26]. The five regions (center, upper, lower, left, and right) were determined based on a pair of crescent-shape furrows.

### Statistical analysis

The morphological parameters, the number and the angle of cell division in Ctrl, *N* RNAi, *CycE* RNAi were conducted in one-way analysis of variance (ANOVA test) followed by the Dunnet test using R console to compare the difference to controls. In this comparison, the number and angle of cell division in each region of *N* RNAi or *CycE* RNAi primordia (a, b, c, d, e in Fig. 3B) were compared to the corresponding region of negative control primordia. For comparing the number and the angle of cell division among regions in each RNAi treatment, multiple comparison analyses were carried out using one-way ANOVA followed by Tukey’s honest significant difference test.

### Gene knockdown via RNAi

We searched the ortholog mRNA sequence from the RNAseq database of *T. dichotomus* (PRJDB6456) using *D. melanogaster* sequences as a query via the tblastn program [27]. Gene knockdown was performed via RNA interference (RNAi) [15]. Briefly, the partial sequence of *Trypoxylus* genes were amplified by a specific primer with an attached T7 sequence. The amplified sequence was cut out from the gel and purified by a gel extraction kit (QIAGEN, Germany). dsRNA was synthesized using the MEGAscript RNA kit (Ambion, Austin, TX, USA) using the purified partial sequence of the *Trypoxylus* gene as a template. Synthesis, purification, and storage of dsRNA were performed according to manufactures protocol. 5 µg of dsRNA was injected into late 3rd instar larvae using a 30 G needle syringe (BD, Japan). The primer sequences were listed in Table S1.

## Acknowledgment

We thank Dr. Shinichi Morita for the helpful discussion about *N* RNAi phenotype and his helpful comments on the manuscript. We also appreciate Drs. Laura Lavine and Mark Lavine for her comments and English correction on the manuscript.

## Author Contributions

Conceptualization: Haruhiko Adachi, Shigeru Kondo and Hiroki Gotoh

Resources: Haruhiko Adachi, Keisuke Matsuda, Hiroki Gotoh

Formal analysis: Haruhiko Adachi

Funding acquisition: Haruhiko Adachi, Teruyuki Niimi, Shigeru Kondo and Hiroki Gotoh

Investigation: Haruhiko Adachi and Hiroki Gotoh

Methodology: Haruhiko Adachi, Keisuke Matsuda and Hiroki Gotoh

Project administration: Shigeru Kondo and Hiroki Gotoh

Supervision: Shigeru Kondo and Hiroki Gotoh

Visualization: Haruhiko Adachi and Hiroki Gotoh

Writing – original draft: Haruhiko Adachi and Hiroki Gotoh

Writing – review & editing: Haruhiko Adachi, Keisuke Matsuda, Teruyuki Niimi, Shigeru Kondo and Hiroki Gotoh

## Supplementary informations

**Figure S1.**
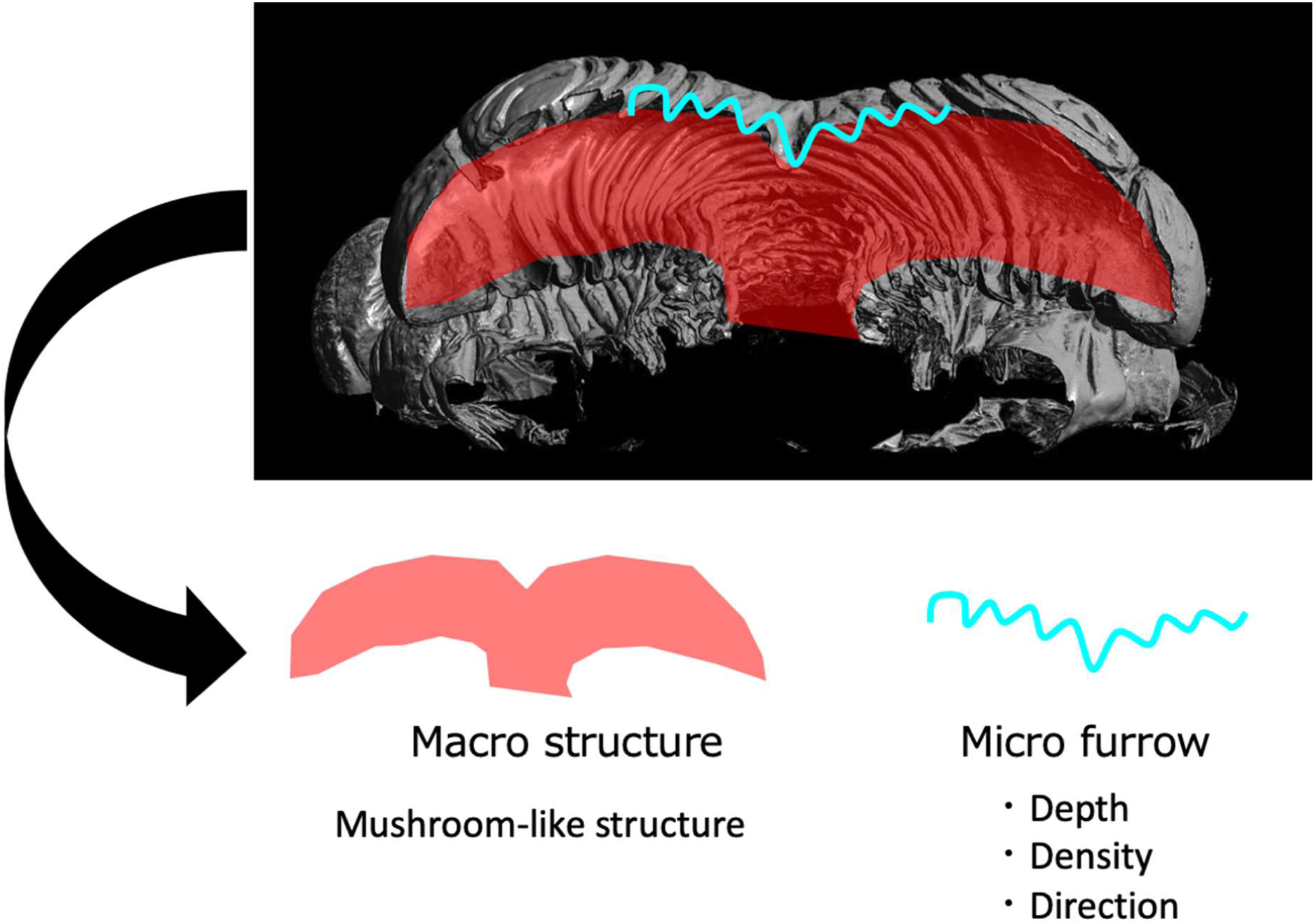
The morphology of the horn primordia. The horn primordia contains a mushroom-like macro structure and surface micro furrows. Micro furrows have depth, density and direction (2D furrow pattern).

**Figure S2.**
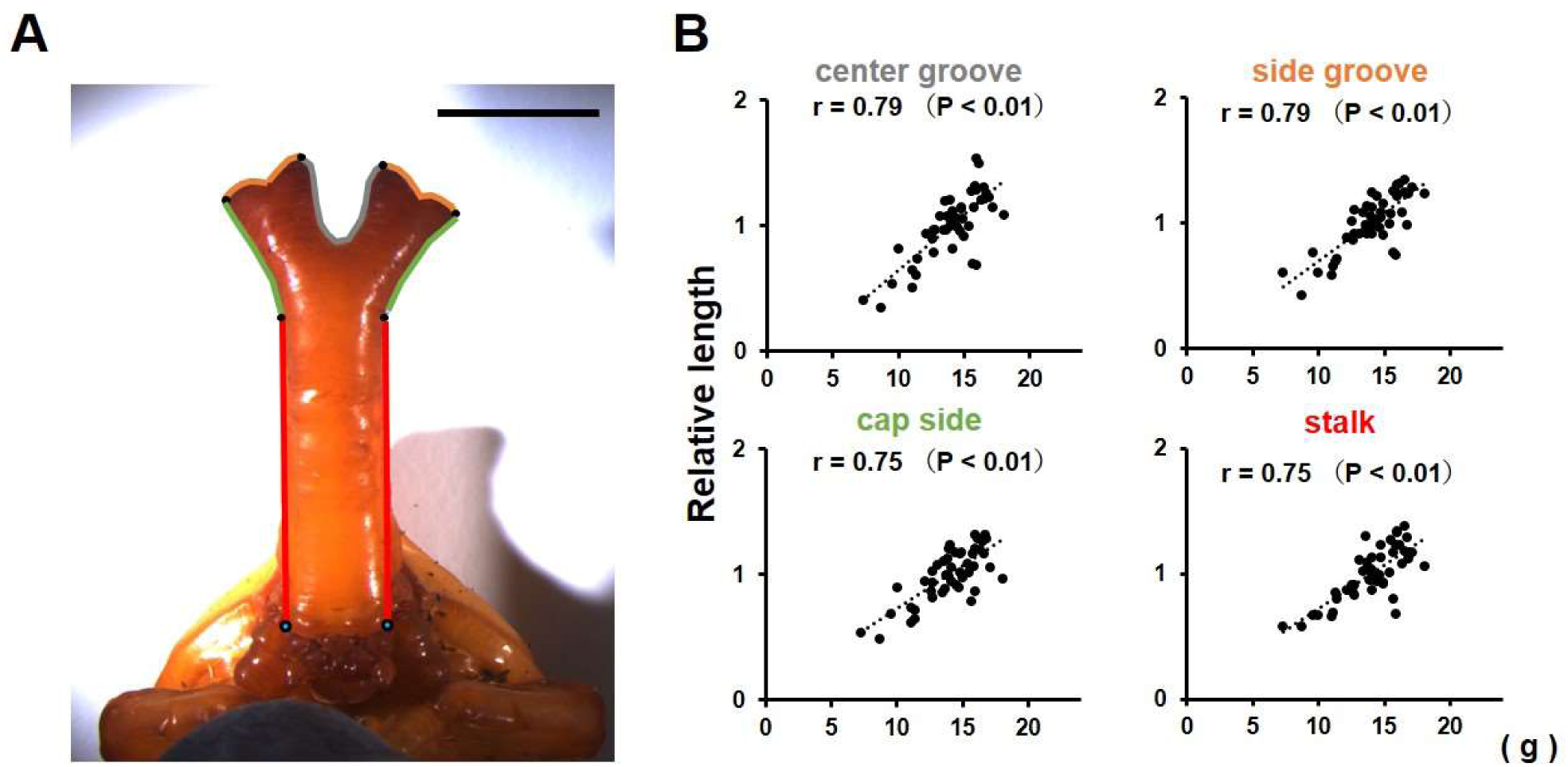
Relationship between body size and the morphology of the pupa. (A) Pupal horn with individual parts outlined. Landmarks were determined by inflection points. Scale bar indicates 10 mm. (B) Relationship between body size and the length of each part of the horn. Measured lengths of each horn part were strongly correlated with body size, and the scaling relationship (the slope) to body size are nearly identical (n = 48).

**Figure S3.**
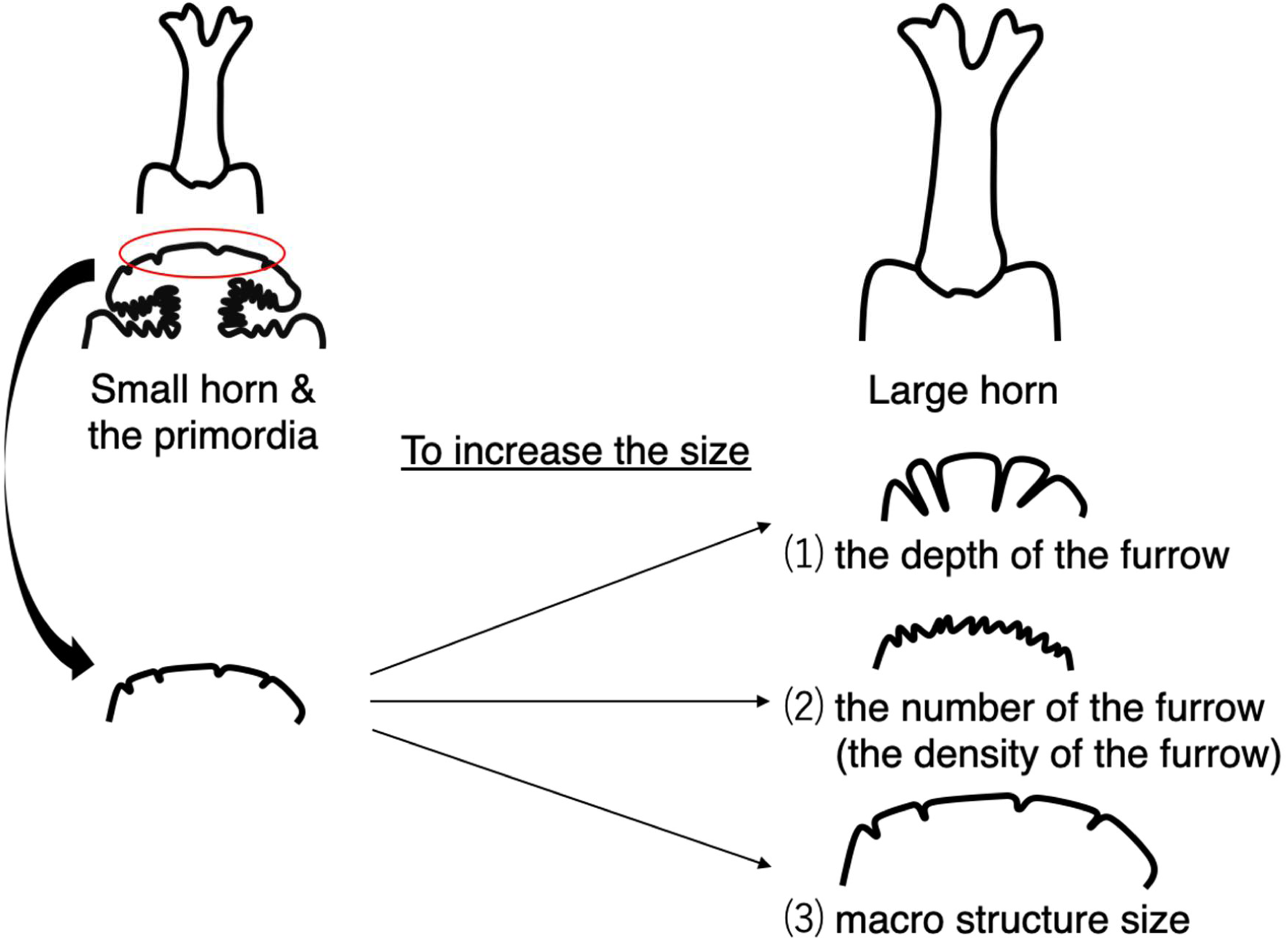
Three way to increase the size of horn via changing morphological parameters of the primordia. There are three way to increase the size of horn via changing morphological parameters of the primordia. First is increasing macro structure size. Second is increasing the number of furrows (in other words, increasing the density of the furrows). Third is increasing the depth of the furrows.

**Figure S4.**
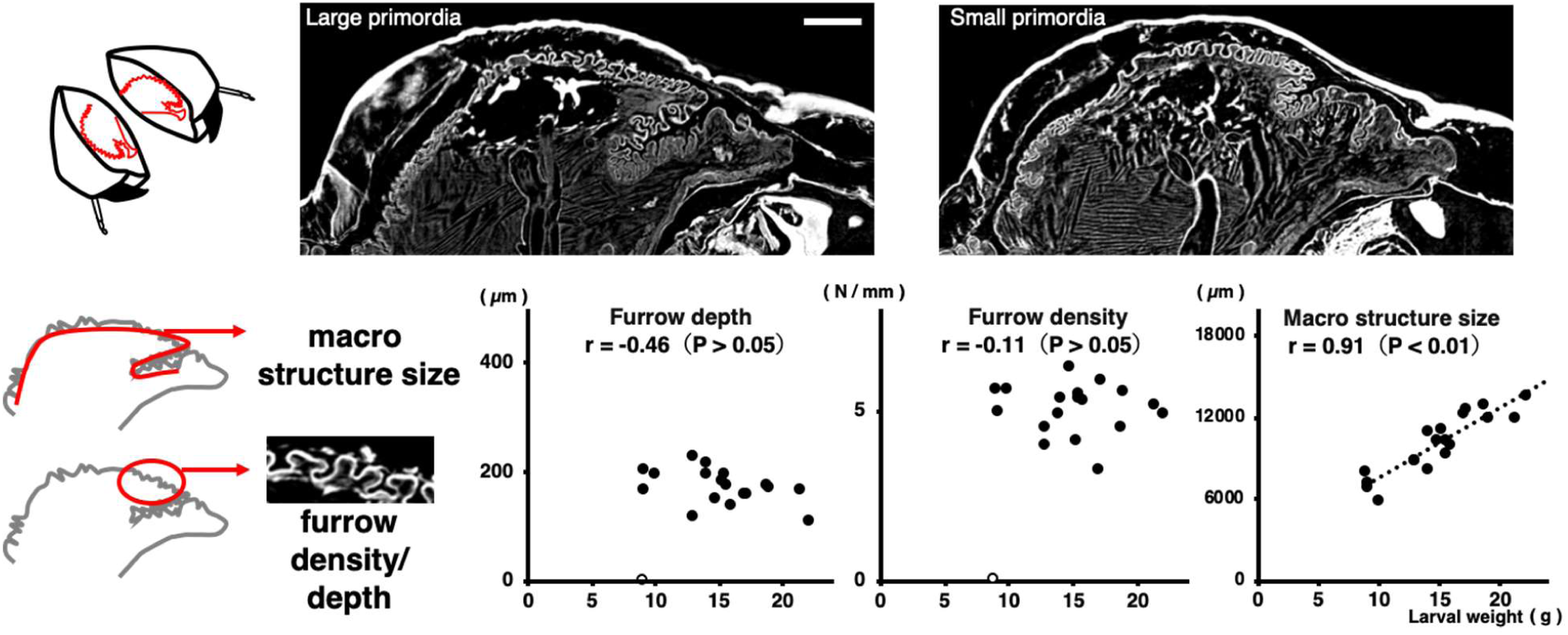
Relationship between body size and the morphology of the primordia (sagittal section) Relationship between body size and the primordial morphological parameters (the macro structure size, the density of the furrows, the depth of the furrows) in the sagittal section. For macro structure size, the correlation coefficient was 0.91 (p < 0.01), for furrow density and depth, the correlation coefficients were not significant. In the smallest beetle (white marker: 8.95 g), obvious furrows could not be detected, so it was excluded from the analysis (n = 19). Scale bar indicates 1 mm.

**Figure S5.**
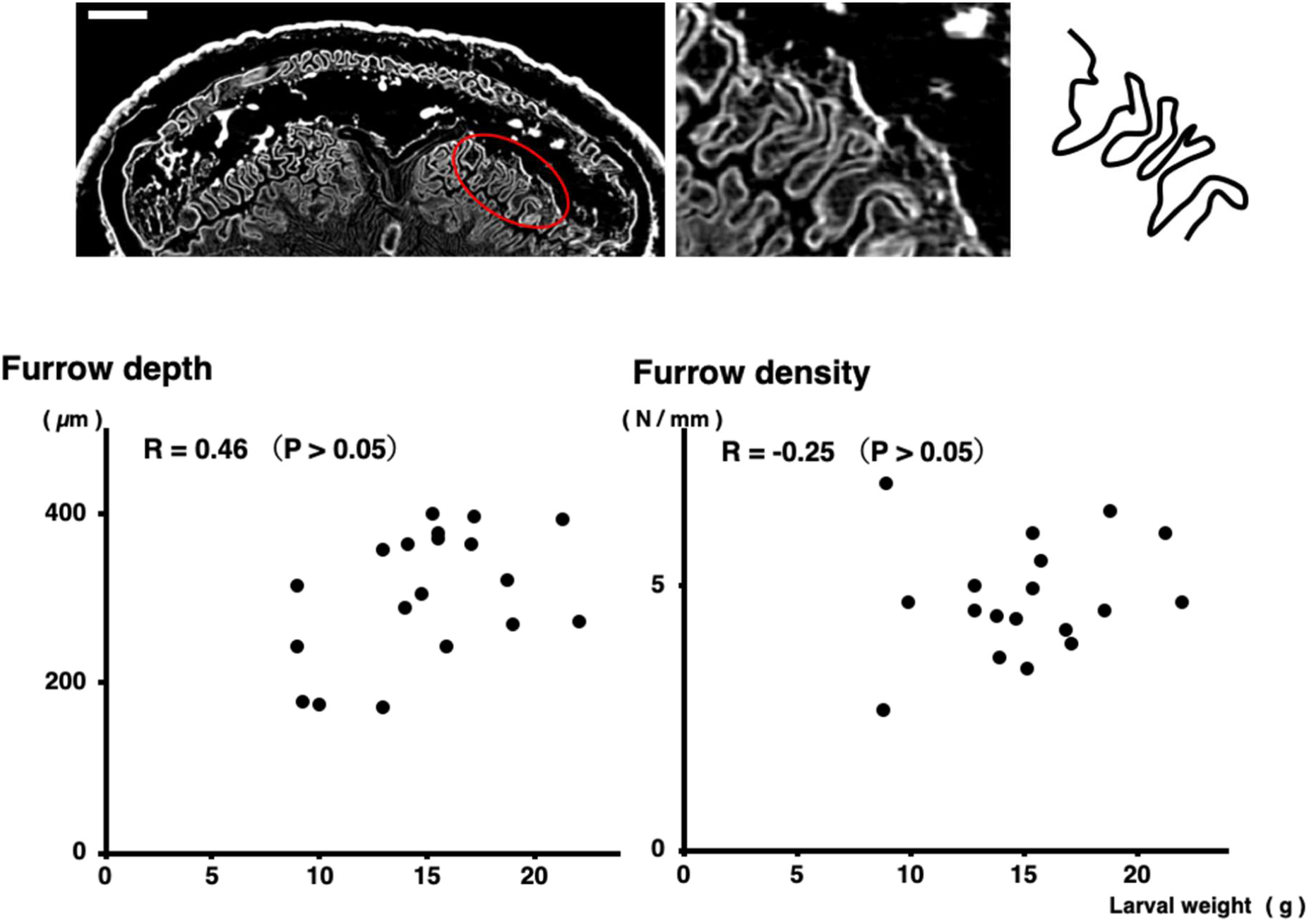
Relationship between body size and the morphology of the primordia (cap bottom region) Relationship between body size and primordia parameters (the macro structure size, the density of the furrows and the depth of the furrows) in the cap bottom region. For furrow density and depth, the correlation coefficients were not significant (n = 19). Scale bar indicates 1 mm.

**Figure S6.**
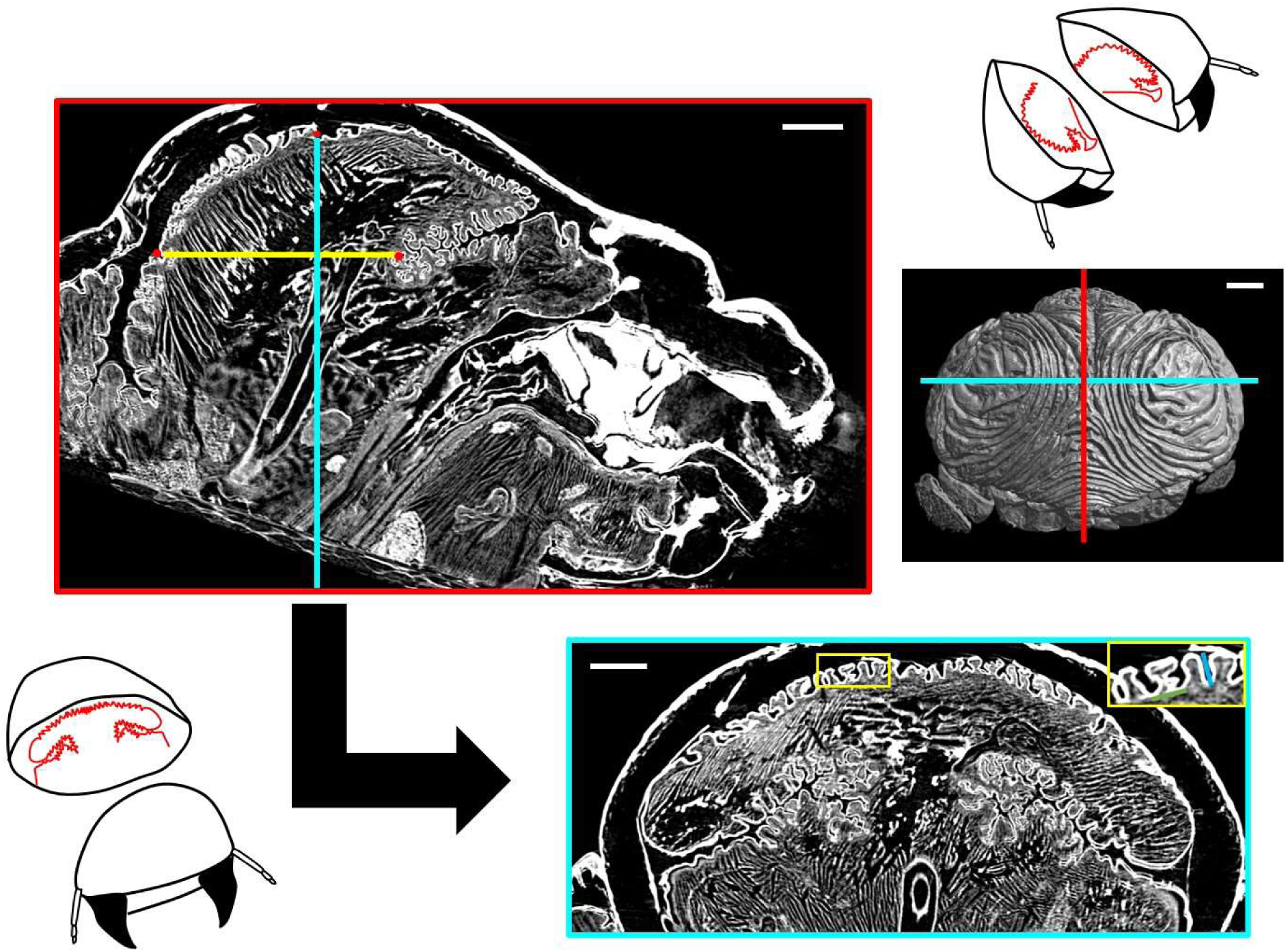
The primordial sectioning point to analyze the morphological parameters. The equal-section images were obtained using landmarks such as the points of changing furrow direction and the inflection point of the macro structure of the primordia. Scale bar indicates 1 mm.

**Figure S7.**
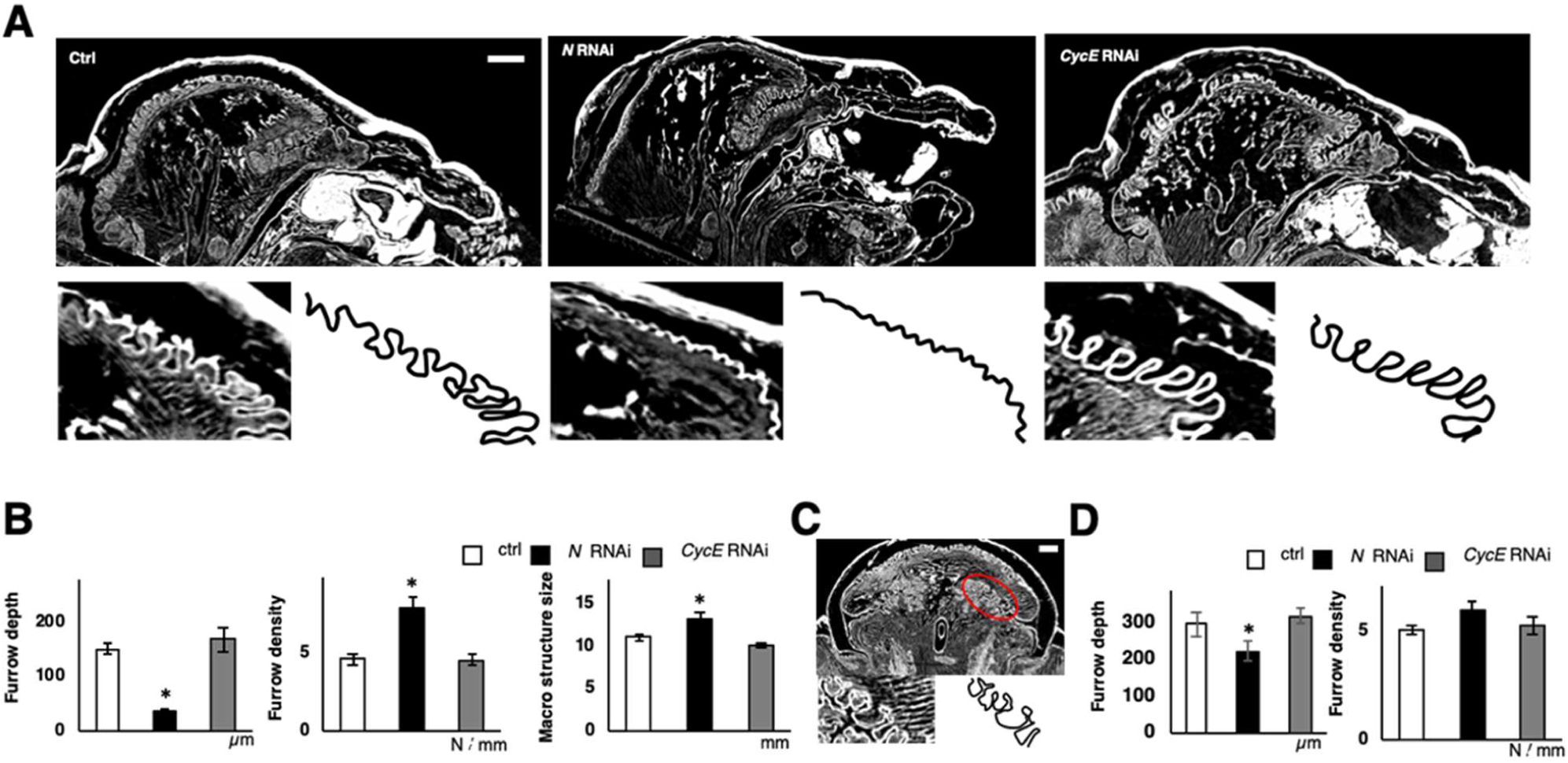
Morphological parameters of the primordia in *N* and *CycE* RNAi. (A) Comparison of sagittal section of the head just before pupation between control and *N* RNAi and *CycE* RNAi. (B) Quantitative data of macro structure size and furrow density and furrow depth from analysis (n = 7, 5, 6 for negative control, *N* RNAi, *CycE* RNAi, respectively). (C) Analysis point of the furrow in cap bottom region. (D) Quantitative data of furrow density and furrow depth in cap bottom region (n = 7, 5, 6 for negative control, *N* RNAi, *CycE* RNAi). Scale bar indicates 1 mm for (A) and (C).

**Figure S8.**
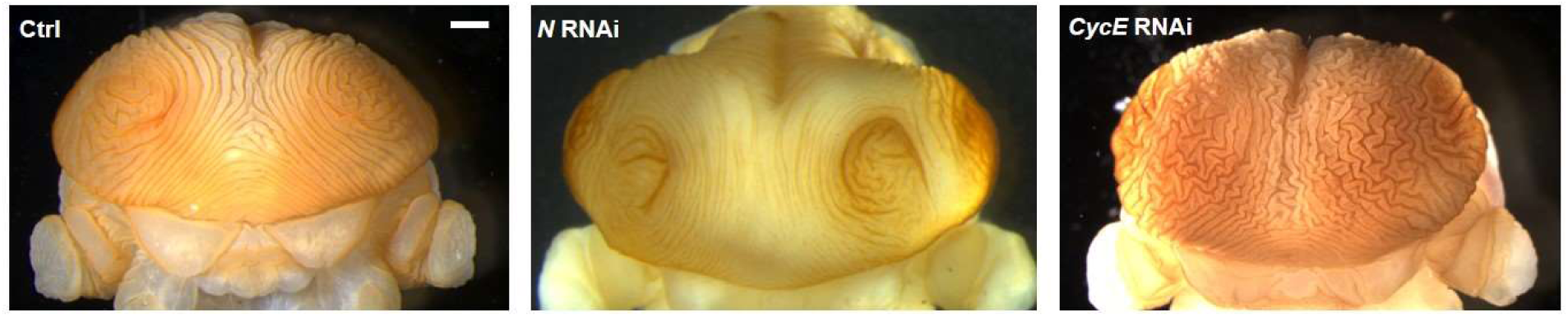
Comparison of 2D furrow pattern between control and *N* and *CycE* RNAi. Comparison of surface 2D furrow pattern between control and *N* and *CycE* RNAi. Two concentric-like furrows are observed in control and *N* RNAi. Zigzag pattern is observed in *CycE* RNAi.

**Figure S9.**
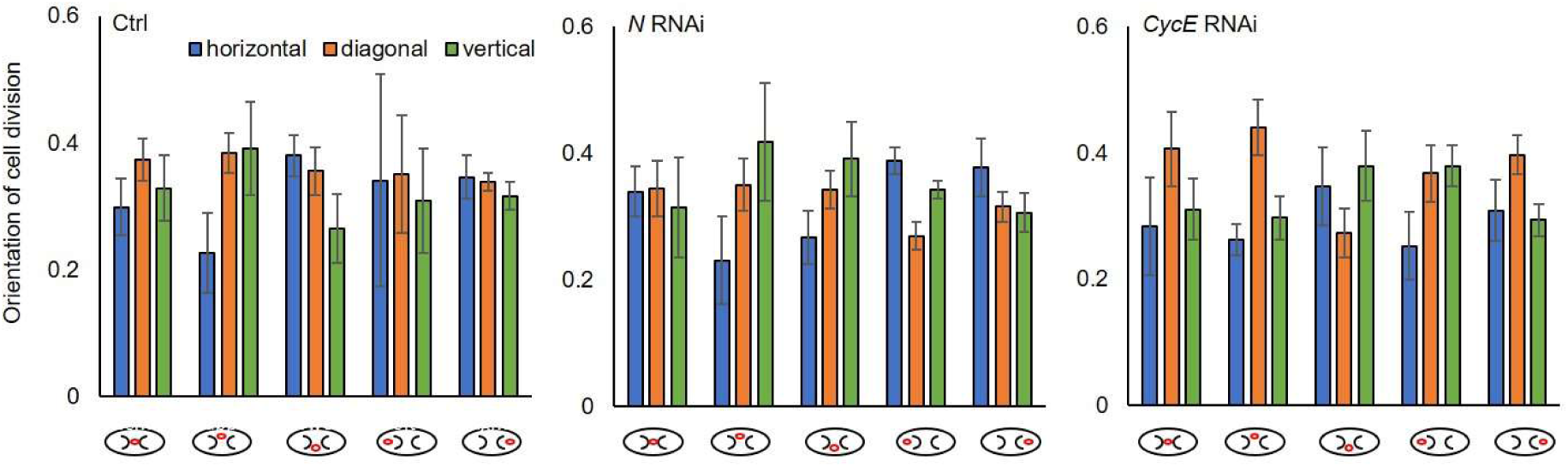
Comparison of cell division angle between control and *N* and *CycE* RNAi. Blue bar shows the ratio of cells divided in horizontal direction (0 < θ ≤ 30). Orange bar shows the ratio of cells divided in diagonal direction (30 < θ ≤ 60). Green bar shows the ratio of cells divided in vertical direction (60 < θ ≤ 90). None of the RNAi treatments showed significant change of cell division orientation for any area.

**Figure S10.**
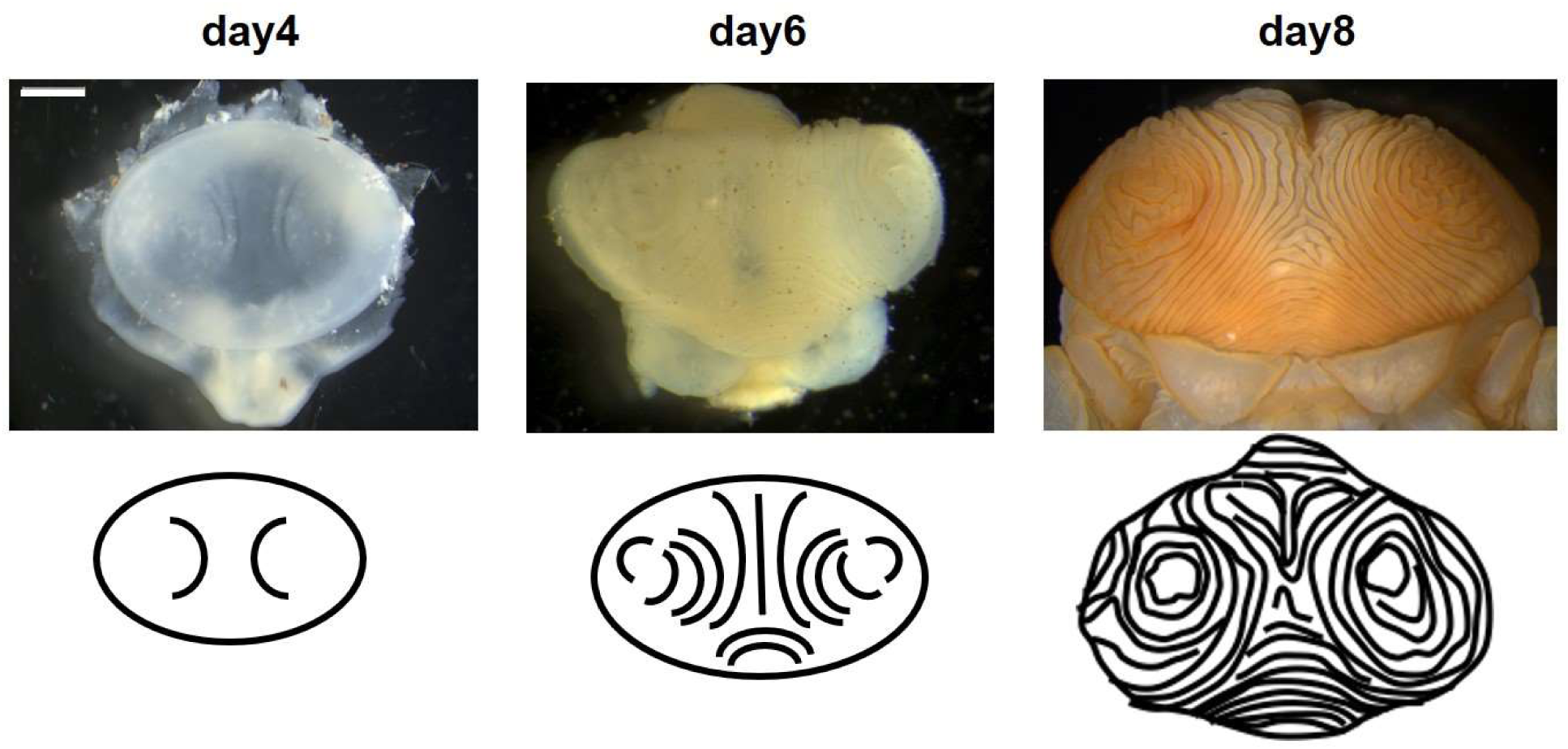
Chronological observation of the primordial furrow formation. A pair of crescent-shape furrows formed in the early stage. We used these furrows as a landmark to investigate cell division patterns across areas. Scale bar indicates 1 mm.

**Table S1:**
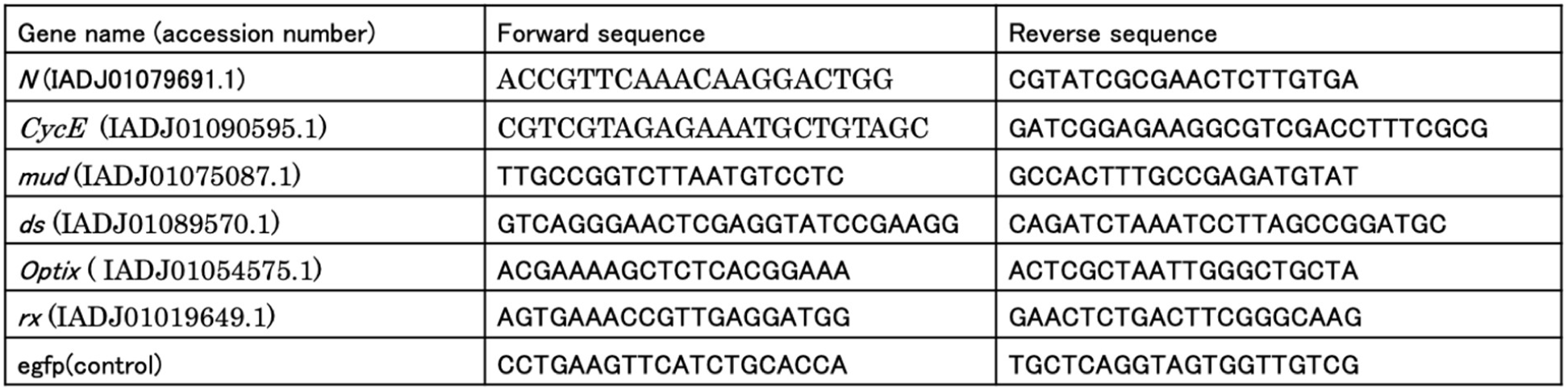
List of primer used for gene amplification as dsRNA synthesis temperate

Supplementary Movie 1 The morphology of the horn primordial

